# Efficacy of heat-killed and formalin-killed vaccines against *Tilapia tilapinevirus* in juvenile Nile tilapia (*Oreochromis niloticus*)

**DOI:** 10.1101/2021.06.03.447010

**Authors:** Thao Thu Mai, Pattanapon Kayansamruaj, Suwimon Taengphu, Saengchan Senapin, Janina Z. Costa, Jorge del-Pozo, Kim D. Thompson, Channarong Rodkhum, Ha Thanh Dong

## Abstract

*Tilapia tilapinevirus* (also known as tilapia lake virus, TiLV) is considered to be a new threat to the global tilapia industry. The objective of this study was to develop simple cell culture-based heat-killed (HKV) and formalin-killed (FKV) vaccines for the prevention of disease caused by TiLV. The fish were immunized with 100 µL of either HKV or FKV by intraperitoneal injection with each vaccine containing 1.8 × 10^6^ TCID_50_ inactivated virus. A booster vaccination was carried out at 21-day post vaccination (dpv) using the same protocol. The fish were then challenged with a lethal dose of TiLV at 28 dpv. The expression of five immune genes (*IgM, IgD, IgT, CD4* and *CD8*) in the head kidney and spleen of experimental fish was assessed at 14 and 21 dpv and again after the booster vaccination at 28 dpv. TiLV-specific IgM responses were measured by ELISA at the same time points. The results showed that both vaccines conferred significant protection, with relative percentage survival (RPS) of 71.3% and 79.6% for HKV and FKV, respectively. Significant up-regulation of *IgM* and *IgT* was observed in the head kidney of fish vaccinated with HKV at 21 dpv, while *IgM, IgD* and *CD4* expression increased in the head kidney of fish receiving FKV at the same time point. After booster vaccination, *IgT* and C*D8* transcripts were significantly increased in the spleen of fish vaccinated with the HKV, but not with FKV. Both vaccines induced a specific IgM response in both serum and mucus. In summary, this study showed that both HKV and FKV are promising injectable vaccines for the prevention of disease caused by TiLV in Nile tilapia.

## 1. Introduction

Tilapia (*Oreochromis* sp.) is the second most farmed fish species worldwide after carps, reaching 6 million tons in 2020 (Fletcher, 2020), equivalent to a value of US$ 7.9 billion (IMARC, 2020). As the demand for animal protein as a food source increases, tilapia has been considered an important freshwater fish for low- and middle-income countries (LMICs) due to its inexpensive price, high adaptability to various environmental conditions and ease to culture (Prabu et al., 2019). Intensification of tilapia farming systems has occurred as a result of this growing demand. This has led to an increased risk of emerging infectious diseases caused by bacteria, viruses, parasites and fungi (Dong et al., 2015; Mesalhy, 2013). In 2013, a new disease with a suspected viral etiology emerged in Ecuador (Ferguson et al., 2014), and was named syncytial hepatitis of tilapia (SHT) based on its characteristic histopathological features. Around the same time, a novel RNA virus causing mass mortalities in tilapia was discovered in Israel, termed tilapia lake virus (TiLV) (Eyngor et al., 2014). Subsequent studies, supported with molecular analysis, revealed that the disease episodes in Ecuador and Israel shared the same causative virus, TiLV (Bacharachet al., 2016b; del-Pozo et al., 2014). The virus has recently been classified as *Tilapia tilapinevirus*, in the *Tilapinevirus* genus, within the *Amnoonviridae* family (Bacharach et al., 2016c). Currently, TiLV has been reported in 16 countries/region worldwide (Jansen et al., 2019; Surachetpong et al., 2020).

Current knowledge indicates that TiLV can infect all stages of fish development, including fertilized eggs, larvae, fry, fingerlings, juveniles and large-size fish (Dong et al., 2017; Senapin et al., 2018) although fingerlings and juveniles appear to be more vulnerable to infection with the virus (Amal et al., 2018; Dong et al., 2017; Ferguson et al., 2014; Surachetpong et al., 2017). Cumulative mortalities of up to 80% have been reported for farmed tilapia in Israel, while in a report from Ecuador the percentage of mortalities appeared to fluctuate from 10-20% up to 80% depending on the fish strain when tilapia fish were transferred to grow out cages, with fish dying within 4-7 days of transfer (Eyngor et al., 2014; Ferguson et al., 2014). The mortality levels caused by TiLV infection in Thailand were also variable, ranging from 20-90% (Dong et al., 2017), and experimental infections also tended to result in high levels of mortality (66-100%) (Behera et al., 2017; Dinh-Hung et al., 2021; Eyngor et al., 2014; Tattiyapong et al., 2017). All of these reports suggest that TiLV is highly virulent and will cause significant mortality loss if introduced to a production site.

Vaccines are an effective way to prevent disease caused by either bacteria or viruses in farmed fish (Evensen, 2016). Currently, the majority of licensed vaccines in aquaculture are inactivated vaccines, which contain either single or combined killed pathogens (Ma et al., 2019; Kayansamruaj et al., 2020), inactivated using either physical (e.g. heat, pH, and ultraviolet) or chemical (e.g. formalin, β-propiolactone, glutaraldehyde) processes (Delrue et al., 2012; Lelie et al., 1987). Ideally, when a vaccine is administered, the fish’s immune response is stimulated to produce of antibodies and an immunologic memory against the pathogen (Secombes & Belmonte, 2016), so that the immune system responses more effectively if the fish should encounter the pathogen at a late date. However, to improve the efficacy of the vaccine, a booster dose(s) is often required in order to obtain high antibody titers against the pathogen (Angelidis, 2006; Bogwald & Dalmo, 2019; Thu Lan et al., 2021). Inactivated vaccines normally stimulate humoral immune responses, involving helper T cells (CD4+ T cells) and antibody-secreting B cells, secreting IgM, IgD or IgT (Smith et al., 2019). The antibodies combat invading pathogens through a variety of mechanisms, including neutralization, phagocytosis, antibody-dependent cellular cytotoxicity, and complement-mediated lysis of pathogens or infected cells (Forthal, 2014). Viral vaccines can also activate cell-mediated immunity, involving cytotoxic T-cells (also known as CD8+ T cells), the function of which is to destroy virus-infected cells (Secombes & Belmonte, 2016; Smith et al., 2019; Somamoto et al., 2002; Toda et al., 2011).

Many vaccines traditionally formulated from inactivated bacteria or viruses, have been licensed and are commercially available for a variety of fish species, mainly salmon, trout and carp (Ma et al., 2019). The few studies that have been reported relating to the development of a vaccine to prevent TiLV infections in tilapia. The first TiLV vaccine was developed in Israel using strains of TiLV that were attenuated by 17-20 subsequent passages in cell culture. The prototype for these vaccines had relative percentage survival (RPS) values of over 50% (Bacharach et al., 2016a). More recently, a cell-culture derived vaccine containing virus inactivated with β-propiolactone and adjuvant Montanide IMS 1312 VG, with a virus titer of 10^8^ 50% tissue culture infectious dose per milliliter (TCID_50_ mL^-1^) was developed in China. The vaccine gave a relatively high level of protection, with the RPS value of 85.7 %. This vaccine was able to induce specific IgM, as well as upregulate a variety of immune genes (Zeng et al., 2021a). In another study, a DNA vaccine consisting of a pVAX1 DNA vector containing the sequence for TiLV’s segment 8, encoding an immunogenic protein VP20, was used for the primary immunization and a recombinant VP20 (rVP20) protein was used as a booster vaccine given at 3-week post-vaccination (wpv). This vaccine combination resulted in a RPS value of 72.5 %, compared to 50 % and 52.5 % respectively for the DNA vaccine or rVP20 alone (Zeng et al., 2021b). In the present study, we investigated whether simple cell culture-based vaccines (water-based with no adjuvant), containing either heat-killed or formalin-killed virus, were able to provoke a specific immune response in vaccinated fish and if the vaccines protected them from TiLV infection.

## 2. Materials and methods

### 2.1. Fish

Juvenile Nile tilapia (*Oreochromis niloticus*) (body weight, 7.3 ± 1.2 g; length, 5.9 ± 1.1) were obtained from a commercial tilapia hatchery with no previous record of TiLV infection. The fish were placed in 100-liter containers at a density of 60 fish per tank at around 28 °C and fed with a commercial diet daily at 3% of body weight for 15 days before performing the vaccination trial. Prior to the experiment, 5 fish were randomly selected to screen for the presence of TiLV using a semi-nested PCR (Taengphu et al., 2020) and bacteria using conventional culture method and found to be negative. Water quality parameters including pH, ammonia, and nitrite concentration was monitored every 3 days using a standard Aqua test kit (Sera, Germany), and water was changed twice per week. The vaccination study was approved by Kasetsart University Institutional Animal Care and Use Committee (ACKU62-FIS-008).

### 2.2. Virus preparation

TiLV strain TH-2018-K was isolated from Nile tilapia during a TiLV outbreak in Thailand in 2018 using E11 cell line following the protocol described previously by Eyngor et al., (2014). The virus was cultured in 75 cm^2^ flasks containing confluent E11 cells and 15 ml of L15 medium at 25°C for 5-7 days or until the cytopathic effect (CPE) of around 80 % was obtained in the cell monolayer. The culture supernatant containing the virus was centrifuged at 4,500 g for 5 min at 4°C (Eppendorf 5810R) and stored at -80°C. The concentration of the virus was determined by calculating the virus titre as 50% tissue culture infective dose per milliliter (TCID_50_ mL^-1^) (Reed & Muench, 1938).

### 2.3. Vaccine preparation

TiLV TH-2018-K (1.8 × 10^7^ TCID_50_ ml^-1^) was used to prepare both HKV and FKV. Viral inactivation was performed at 60 °C for 2, 2.5, and 3 h or with formalin (QReC) at a final concentration of 0.002%, 0.004%, 0.006%, 0.008% and 0.01% for 24 h at 25°C. Viral infectivity was then checked on E11 cells. Successful inactivation of the virus was confirmed by the absence of a cytopathic effect (CPE) after 7 days with all inactivation conditions tested (Table S1). Subsequently, inactivation of the virus was performed at 60°C for 2.5 h for HKV, while incubation of 0.006% formalin at 25°C for 24 h was used for FKV. The inactivated viral solutions were used as vaccine preparations and were not adjuvanted. These were stored at 4°C until used.

### 2.4. Immunization, sampling and challenge test

Before immunization, 6 fish were chosen randomly from the fish population for blood and mucus sampling. The vaccine study comprised of three groups (HKV, FKV and control). Each group consisted of two 100-L replicate tanks with 25 fish each. Prior to vaccination, fish were anaesthetized using clove oil (100 ppm). Fish in the vaccine groups were immunized with either HKV or FKV by intraperitoneal (IP) injection with 100 µL of vaccine using a 28G × 13 mm needle. Booster immunization was carried out at 21 dpv with the same dose of vaccine (Table 2). Fish in the control group were treated the same, except L15 medium was used in place of the virus solution. Three fish from each tank were randomly collected at 14, 21 and 28 dpv for blood, mucus and tissue sampling (6 biological replicates per treatment). Before sampling, fish were anaesthetized with clove oil at 100 ppm. Mucus samples were collected from each fish by placing the fish into a plastic bag containing 1 mL phosphate-buffered saline (PBS, 137 mM NaCl, 2.7 mM KCl, 10 mM Na_2_HPO_4_, and 1.8 mM KH_2_PO_4_) followed by gentle rubbing for 30s. These were then centrifuged at 4,000 g for 10 min. The mucus supernatant samples were collected and stored at -20°C until used. Blood (∼ 200 µL) was withdrawn from caudal vessel using a 25G × 16 mm needle and allowed to clot for 2 h at 4°C. Serum was collected after centrifugation the blood at 4,000 g for 10 min (Thermo Scientific, UK) and then stored at -20°C. Tissues (head kidney and spleen) were collected, immediately placed in Trizol solution (Invitrogen, UK), and kept at -20°C until RNA extraction. For the challenge test, a viral stock of TiLV strain TH-2018-K (1.8 × 10^7^ TCID_50_ mL^-1^) was diluted 2 times with sterile distilled water. Each fish was injected IP with 0.1 mL of the diluted TiLV solution (9 × 10^5^ TCID_50_ fish^-1^) at 28 dpv, and mortalities were monitored daily for 21 days. Representative dead fish from each group were subjected for TiLV diagnosis using an in-house RT-qPCR (Taengphu et al., submitted).

### 2.5. Immune-related gene expression by RT-qPCR

RNA was extracted using Trizol (Invitrogen, UK) following the protocol recommended by the manufacturer. Genomic DNA contamination was removed using DNase I (Ambion, UK) according to the manufacturer’s instructions. After DNase I treatment, RNA samples were re-purified using an equal volume of acid phenol:chloroform (5:1, pH 4.7) (Green & Sambrook, 2019) before checking quality and quantity of extracted RNA with Nanodrop ND-1000 Spectrophotometer (Thermo Scientific, UK). DNA contamination in the treated RNA samples was assessed by performing a qPCR cycling with tilapia elongation factor 1α (*EF-1α*) primers using No-RT master mix (absence of reverse transcriptase enzyme provided in iScript™ Reverse Transcription kit, Bio-Rad, USA). The cDNA synthesis (20 µL reactions) was performed using an iScript™ Reverse Transcription Supermix (Bio-Rad, USA) containing 100 ng RNA and incubated at 25°C for 5 min for priming, followed by 46°C for 20 min for reverse transcription and then 95°C for 1 min for inactivation of the reverse transcriptase. Immune-related gene expression in the head kidney and spleen were analyzed using a quantitative real-time PCR, with specific primers as listed in Table 1 and iTaq Universal SYBR Supermix (Bio-Rad, USA). The 10 µL reaction consisted of 5.0 µL 2X Supermix, 0.5 µL forward and reverse primers (10 µM each), 1.0 µL cDNA and 3.0 µL distilled water. The reaction consisted of an initial activation at 95°C for 2 min, followed by 40 amplification cycles of denaturation at 95°C for 30 s, annealing at the optimal temperature of each primer pair (as shown in Table 1), and extension at 72°C for 30 s. Gene expression data for the immune-related genes of vaccinated and control fish were normalized with that of *EF-1α* gene amplification using the 2^-ΔΔCt^ method (Livak & Schmittgen, 2001).

**Table 1.**
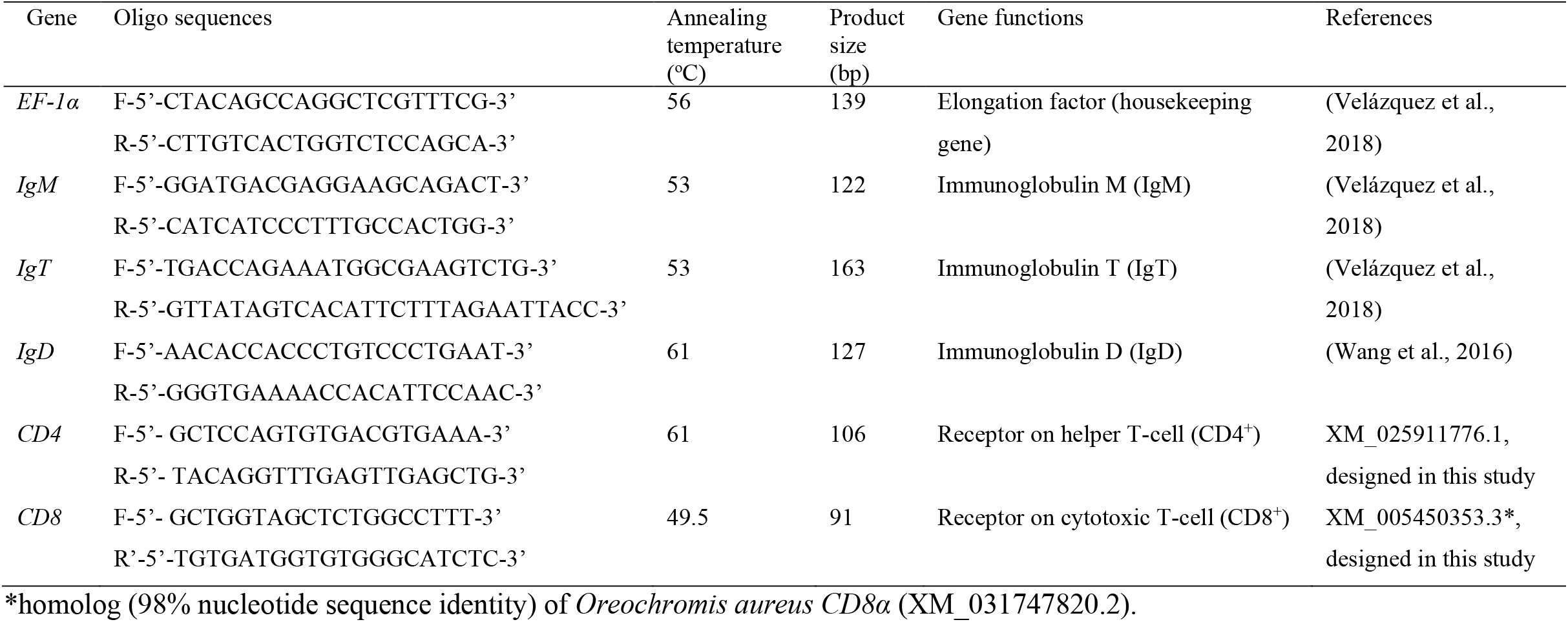
Details of primers used for immune-related gene expression in this study.

### 2.6. Measurement of antibody response by ELISA

Polystyrene 96 well ELISA plates were coated with 0.01% poly-L-lysine solution for 1 h. The plates were then rinsed 3 times with low salt wash buffer (LSWB, 2 mM Tris; 38 mM NaCl; 0.005% Tween 20, pH 7.3) before the addition of 100 µL of either heat- or formalin-inactivated TiLV (1.8 × 10^7^ TCID_50_ mL^-1^) overnight at 4°C. The plates were washes 3 times with LSWB, followed by a blocking step with PBS + 1% bovine serum albumin (BSA, Sigma) for 2 h at room temperature (around 28°C). Then, 100 µL mucus (undiluted) or sera (diluted 1:512 in PBS) were added to each well and incubated overnight at 4°C. The following day, the plates were washed 5 times with high salt wash buffer (HSWB, 2 mM Tris; 50 M NaCl; 0.01% Tween 20, pH 7.7) and incubated with anti-tilapia IgM (Soonthonsrima et al., 2019) diluted at the ratio 1:200 in PBS + 1% BSA for 2 h at around 28°C. The plates were then washed 5 times with HSWB followed by incubation of goat anti-mouse antibody (Merck, Germany) conjugated with HRP (diluted 1:3000 in LSWB + 1% BSA) for 1 h at around 28°C. The plates were finally washed 5 times with HSWB before adding 100 µL of TMB (Merck, Germany) to each well. Color was developed in the dark for 5-10 min before adding 50 µL of 2 M H_2_SO_4_ stop solution (Merck, Germany). Optical density was read at wavelength 450 nm using the microplate reader (SpectraMax ID3, USA).

### 2.7. Statistical analysis

GraphPad Prism 6 was used to generate the graphs. Kaplan-Meier analysis was performed and the log-rank test was used to compare the survival curves between vaccinated and control groups. The relative percentage survival (RPS) was calculated using following equation:

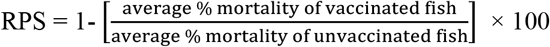

The differences in relative fold change of immune-related gene expression and specific antibody IgM level were compared using two-way ANOVA followed by the LSD post hoc test. The differences are considered at different levels of significance *p <0*.*05, p<0*.*01, p<0*.*001* and *p<0*.*0001*.

## 3. Results

### 3.1. Efficacy of vaccine

In the challenge experiment, the first mortality occurred at 3-day post challenge (dpc) in the non-vaccinated group (control) and at 5 and 7 dpc in the HKV and FKV groups, respectively (Fig. 1). Mortalities continued until 13-15 dpc. Moribund fish showed gross signs of TiLV infection including abdominal distension, skin erosion, exophthalmos, fin rot, gill pallor and pale liver (Fig. S1). The dead fish from each group were tested positive for TiLV by RT-qPCR. The survival rates were 81.3 ± 0.0 % and 86.3 ± 0.0% for HKV and FKV groups, respectively, compared to 28.13 ± 30.9% for the control (*p<0*.*0001*). The survival percentage were analysed using Kaplan-Meier curves with the log rank test (Fig. 1). Average RPS values were 71.3 % for the HKV and 79.6 % for the FKV vaccine (Table 2).

**Figure 1.**
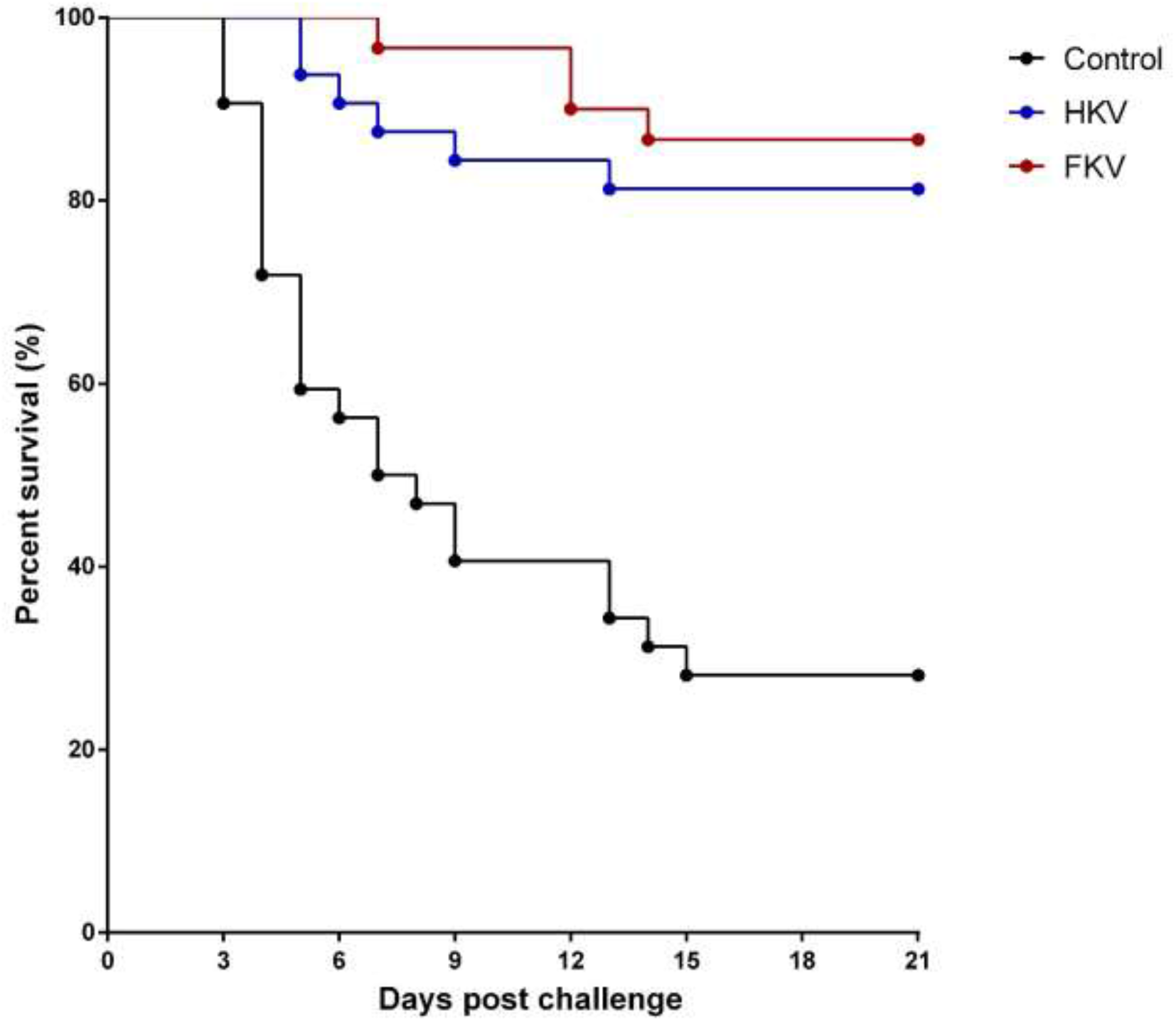
Average percent survival of heat-killed and formaldehyde-killed vaccinated groups (HKV vs. FKV) compared to the non-vaccinated group (Control) during 21 days post challenge. with TiLV (strain TH-2018-K). Statistical analysis of cumulative survival between both vaccinated groups and the control were analyzed using Kaplan-Meier curve with log-rank test (*p < 0*.*0001*).

**Table 2.** Details of experimental groups and challenge results

| Treatment | Administration route | No of fish challenged | Primary vaccination (TCID<sub>50</sub> fish<sup>-1</sup>) | Booster vaccination (TCID<sub>50</sub> fish<sup>-1</sup>) | Challenge (TCID<sub>50</sub> fish<sup>-1</sup>) | Survival rate (%) | RPS (%) | Significant level (compared to control) |
| --- | --- | --- | --- | --- | --- | --- | --- | --- |
|  |  |  | Day 0 | Day 21 | Day 28 |  |  |  |
| Control (L15 media) | IP | 16 (× 2 rep.) | 0 | 0 | $9 \times 10^5$ | $28.13 \pm 30.9$ | NA | |
| HKV | IP | 16 (× 2 rep.) | $1.8 \times 10^6$ | $1.8 \times 10^6$ | $9 \times 10^5$ | $81.3 \pm 0.0$ | 71.3 | $p < 0.0001$ |
| FKV | IP | 15 (× 2 rep.) | $1.8 \times 10^6$ | $1.8 \times 10^6$ | $9 \times 10^5$ | $86.3 \pm 0.0$ | 79.6 | $p < 0.0001$ |

HKV, heat-killed vaccine group; FKV, formalin killed vaccine group; rep, replicate; NA, not applicable; IP, intraperitoneal injection

### 3.2. Immune-related gene expression

The relative fold changes of five immune genes (*IgM, IgT, IgD, CD4, CD8*) were compared to that of the control group (Fig. 2). In the head kidney, a non-significant increase of *IgM* mRNA relative to the control was noted at 14 dpv, which was followed by significant increase relative to the control at 21 dpv for both HKV and FKV groups (Fig. 2A, *p<0*.*05*). A similar trend was observed for *IgT* at 14 dpv for both vaccine groups, which was followed by significantly higher expression levels at 21 dpv for the HKV group only (Fig. 2B, *p<0*.*05*). Regarding mRNA levels of *IgD*, there was significant up-regulation of *IgD* in the FKV group only at 21 dpv (Fig. 2C, *p<0*.*01*). The *CD4* gene was significantly upregulated at 14 dpv in the HKV only (Fig. 2D, *p<0*.*05*) and at 21 dpv in the FKV (*p < 0*.*001*). No statistical difference was observed in *CD8* expression between the vaccinated and control groups at the time point examined (Fig. 2E).

**Figure 2.**
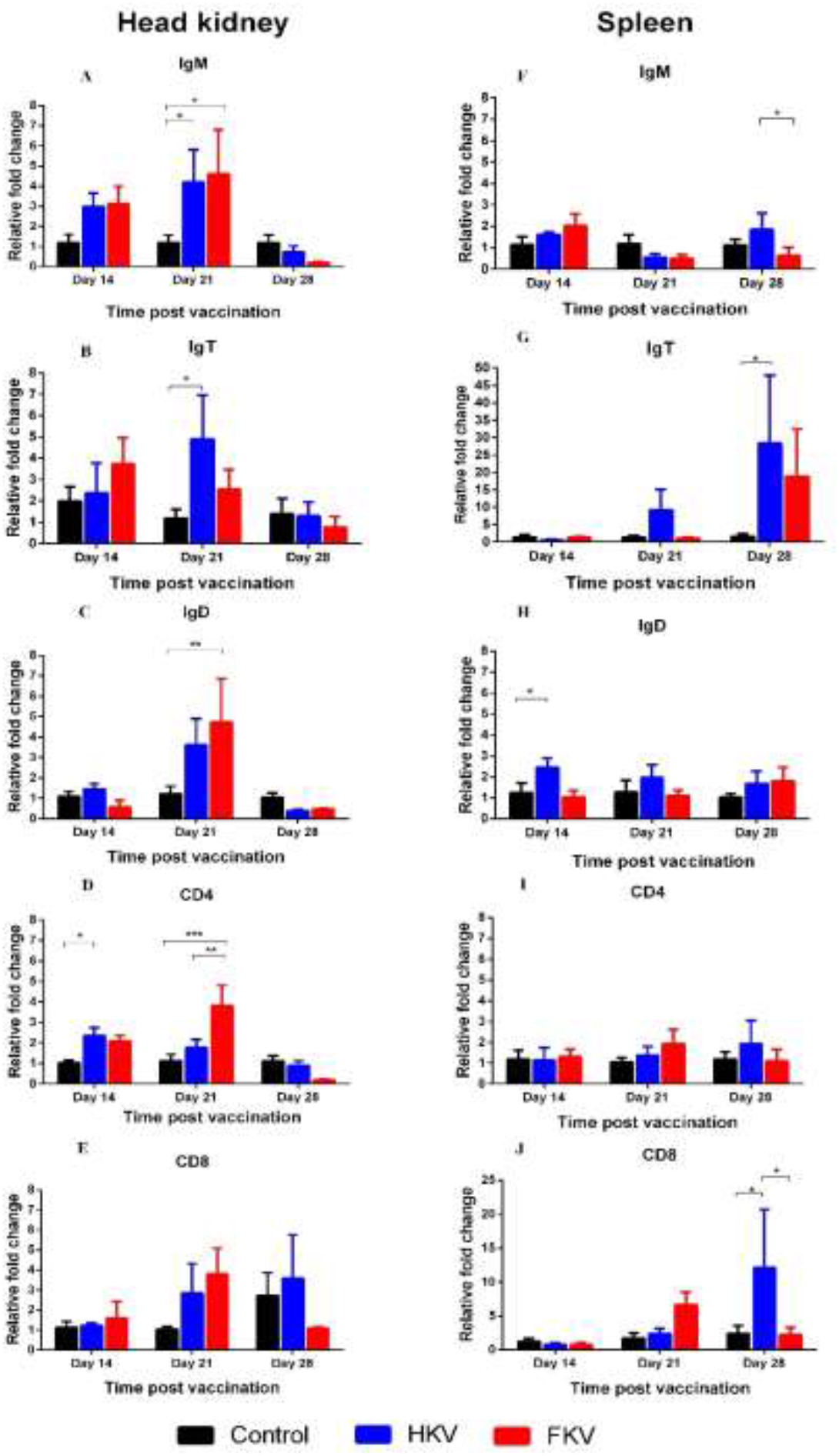
Fold change in gene expressions between non-vaccinated and vaccinated fish at 14, 21 and 28 - day post vaccination. Data are presented as the mean ± SE (n=6). Control, non-vaccinated group; HKV, heat-killed vaccine group; FKV, formalin-killed vaccine group. Asterisks show significant levels between groups. * *p<0*.*05*, ** *p< 0*.*01*, *** *p<0*.*001*

In the spleen, non-significant, relative up-regulation of *IgM* expression was noted in both HKV and FKV groups compared to the control at 14 dpv. (Fig. 2F). There was a slight increase of *IgM* mRNA level relative to the control in the HKV group after booster (28 dpv), which were not significant. Also at 28 dpv, *IgT* expression was over 25 times higher in the HKV group (*p<0*.*05*) and almost 20 times higher in the FKV group (Fig. 2G). A slight significant increase in *IgD* expression was seen in the HKV group at 14-dpv (Fig. 2H, *p<0*.*05*). No significant increase of *CD4* expression was found at any time point (Fig. 2I); meanwhile, an approximately tenfold increase of *CD8* expression was observed at 28 dpv in the HKV group (Fig. 2J, *p<0*.*05*).

### 3.3. Detection of antibody IgM against TiLV in serum and mucus

Systemic TiLV-specific antibody IgM (anti-TiLV IgM) levels pre-vaccination (0 dpv) and at 14, 21 and 28 dpv, as indicated by optical density (OD) at 450 nm, were determined by ELISA (Fig. 3A). Before immunization, the average OD value of the fish sera was 0.096 ± 0.009. The OD readings for HKV, FKV and control groups were 0.254 ± 0.053, 0.363 ± 0.09 and 0.096 ± 0.015 at 14 dpv, respectively. The OD values showed an increase in antibody levels in both groups of vaccinated fish, but were only statistically different in the FKV group (*p<0*.*01*). A slight decrease was seen in OD readings at 3 wpv in both the HKV and FKV groups relative to the control group (0.249 ± 0.049, 0.317 ± 0.043 and 0.128 ± 0.017, respectively). One week after the booster vaccination at 28 dpv, the anti-TiLV IgM levels had increased considerably in both the HKV (*p<0*.*001*) and the FKV (*p<0*.*0001*) groups, reaching the highest values obtained between the different sampling points, compared to that of the non-vaccinated group (average OD readings were 0.438 ± 0.127, 0.483 ± 0.088, and 0.081 ± 0.01 respectively) (Fig. 3A).

**Figure 3.**
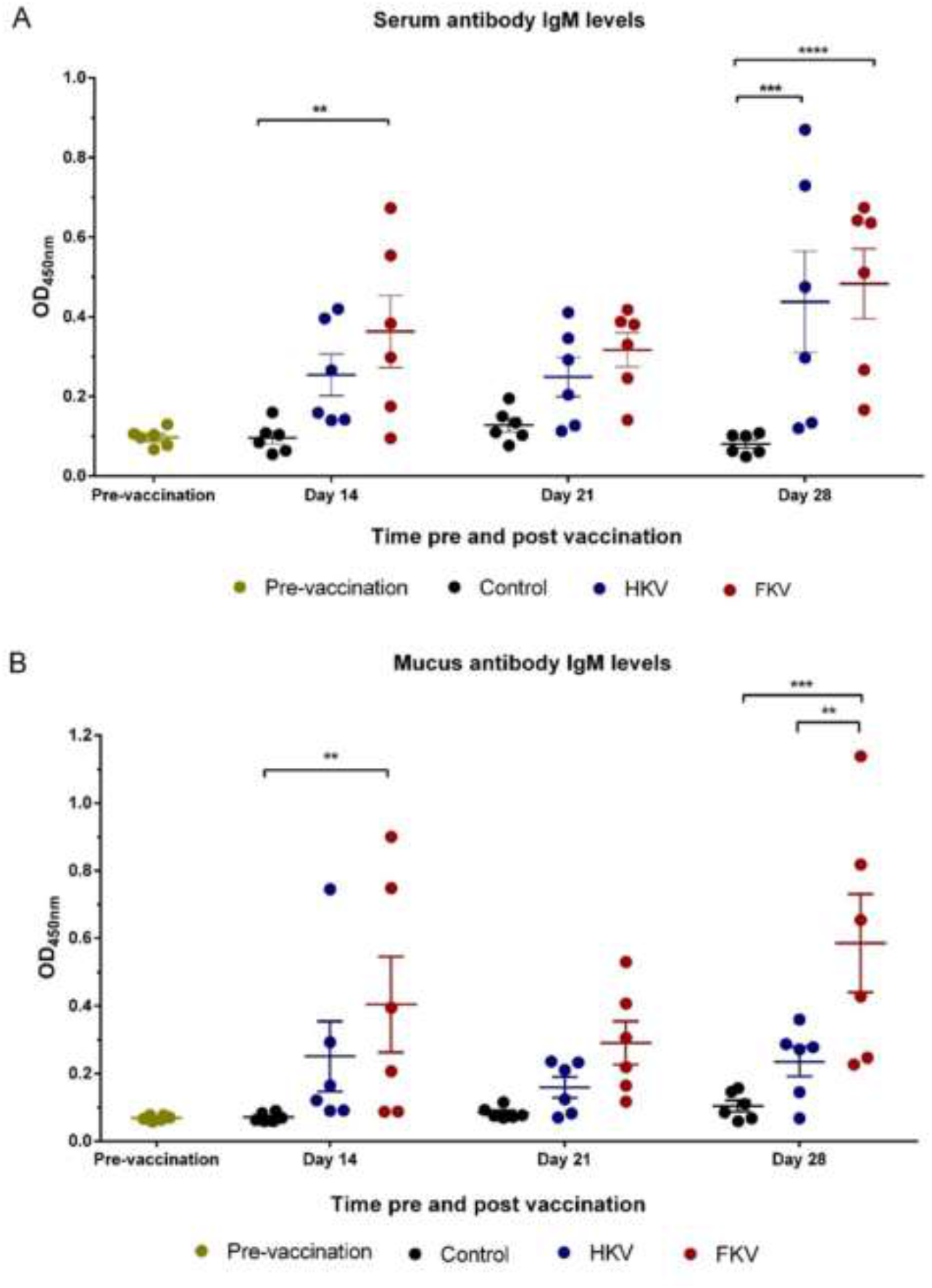
Optical Density (OD) at 540 nm for IgM levels against TiLV in fish sera (diluted 1:512) (A) and mucus (undiluted) (B). Data are presented as the mean ± SE (n=6). Control, non-vaccinated group; HKV, heat-killed vaccine group; FKV, formalin-killed vaccine group. Asterisks indicate significant levels between groups. * *p<0*.*05*, ** *p< 0*.*01*, *** *p<0*.*001*, **** *p<0*.*0001*.

A similar pattern was observed with the mucosal anti-TiLV IgM response (Fig. 3B). Before vaccination, the average OD value of fish mucus was 0.068 ± 0.003. At 14 dpv, the TiLV-specific antibody IgM rose in both of the vaccinated groups, HKV and FKV, compared to the non-vaccinated group (0.251 ± 0.104, 0.404 ± 0.142, and 0.07 ± 0.005, respectively), but a significant difference was only noted for the FKV group (*p<0*.*01*). At 3 wpv, the antibody levels were not significantly differ between groups, with OD values of 0.159 ± 0.031 (HKV), 0.290 ± 0.064 (FKV), and 0.083 ± 0.007 (control) being recorded. At 4 wpv (after administering the booster vaccination), a considerable increase in anti-TiLV IgM levels was seen in the mucus of the FKV group (*p<0*.*001*) (0.585 ± 0.145), whereas the increase measured in HKV fish (0.235 ± 0.044) was not statically different to that of the control group (0.107 ± 0.018). No significant changes in average OD readings were seen between the non-vaccinated group and pre-immunized fish in either sera or mucus (Fig. 3 A-B).

## 4. Discussion

### 4.1. Both simple HKV and FKV were effective in protecting tilapia from TiLV infection

Although many different types of vaccines have been developed for aquaculture in recent years, whole-cell inactivated vaccines remain the major type of vaccine licensed for use by the aquaculture industry (Kayansamruaj et al., 2020; Ma et al., 2019). They are safe, relatively simple to produce, and are affordable for farmers, especially for species that are intensively cultured, but low in price like tilapia in LMICs. In this study, we prepared two versions of simple water-based inactivated vaccine (HKV and FKV) for TiLV and assessed the ability of both to protective tilapia against the virus. Both HKV and FKV were able to confer relatively high levels of protection (RPS, 71.3% vs. 79.6%) in vaccinated fish. Differences in methods used to inactivate the virus, vaccine formulation, viral strains, antigen concentration, route of vaccine administration and the population of fish can all contribute to the level of protection obtained from a vaccine (Table 3). Despite this, vaccination is still considered as a promising strategy to protect tilapia from TiLV infection, although the design of the vaccine should be carefully considered to optimize the level of protection obtained. Other inactivated vaccines have shown relatively high levels of protection in fish. For example, other formalin-killed vaccines resulted in RPS values of 79%, 81.9% and 74% for infectious hematopoietic necrosis virus in rainbow trout (*Oncorhynchus mykiss*) (Tang et al., 2016), *Betanodavirus* in European sea bass (*Dicentrarchus labrax*) (Nuñez-Ortiz et al., 2016), and scale drop disease virus (SDDV) in Asian sea bass (*Lates calcarifer*) (de Groof et al., 2015), respectively. In addition, a heat-killed *Aeromonas hydrophila* vaccine gave 84% protection in rainbow trout (Dehghani et al., 2012). Although the efficacy of these and the current vaccines were not tested against heterologous strains of TiLV, the high level of protection elicited against the homologous strain suggests that autogenous inactivated vaccines may be effective as an emergency vaccine to reduce the risk of production losses in affected tilapia farms.

**Table 3.**
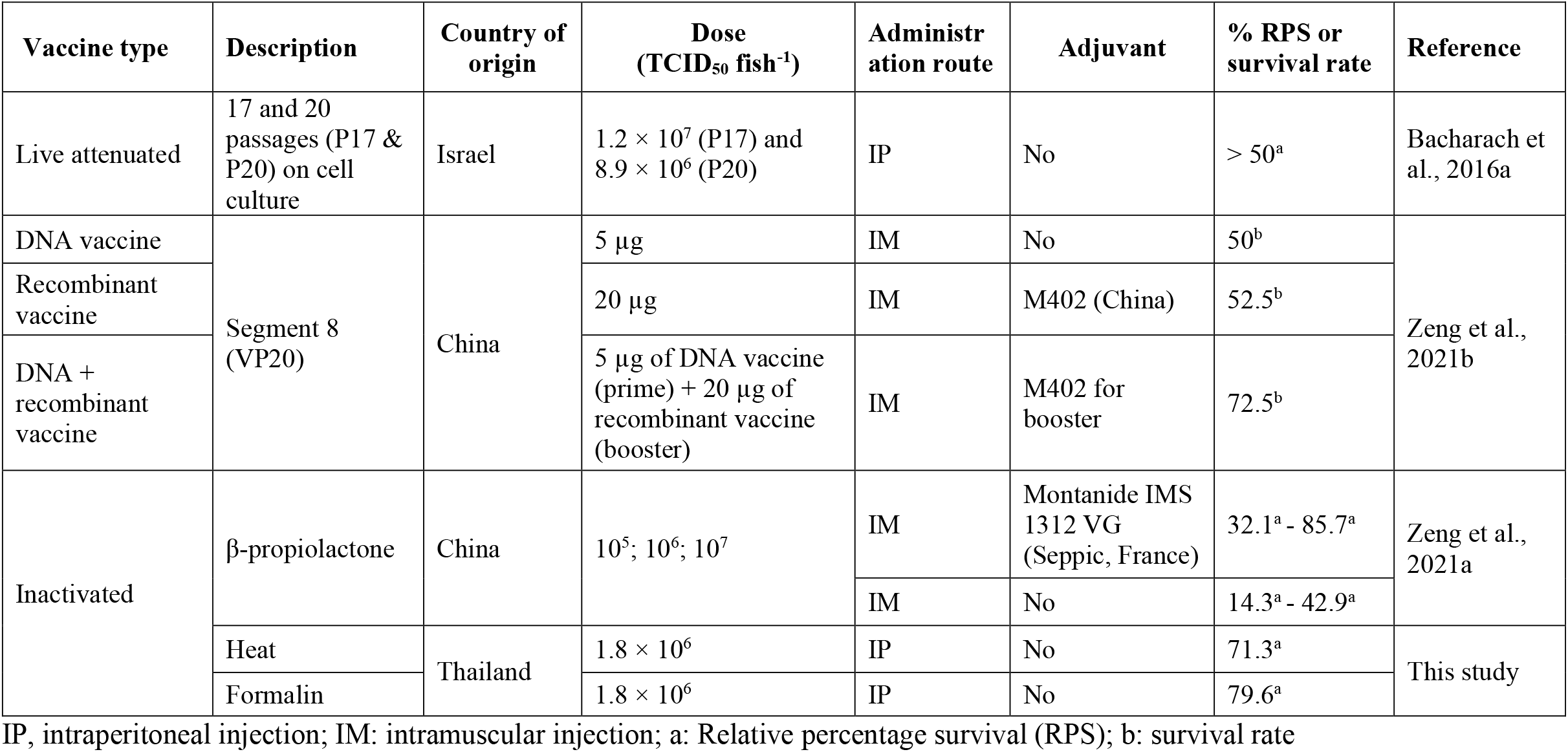
Summary of TiLV vaccines and their efficacy

| Vaccine type | Description | Country of origin | Dose (TCID<sub>50</sub> fish<sup>-1</sup>) | Administration route | Adjuvant | % RPS or survival rate | Reference |
| --- | --- | --- | --- | --- | --- | --- | --- |
| Live attenuated | 17 and 20 passages (P17 & P20) on cell culture | Israel | $1.2 \times 10^7$ (P17) and $8.9 \times 10^6$ (P20) | IP | No | > 50<sup>a</sup> | Bacharach et al., 2016a |
| DNA vaccine | Segment 8 (VP20) | China | 5 µg | IM | No | 50<sup>b</sup> | Zeng et al., 2021b |
| Recombinant vaccine |  |  | 20 µg | IM | M402 (China) | 52.5<sup>b</sup> |  |
| DNA + recombinant vaccine |  |  | 5 µg of DNA vaccine (prime) + 20 µg of recombinant vaccine (booster) | IM | M402 for booster | 72.5<sup>b</sup> |  |
| Inactivated | β-propiolactone | China | $10^5$; $10^6$; $10^7$ | IM | Montanide IMS 1312 VG (Seppic, France) | 32.1<sup>a</sup> - 85.7<sup>a</sup> | Zeng et al., 2021a |
|  |  |  |  | IM | No | 14.3<sup>a</sup> - 42.9<sup>a</sup> |  |
| | Heat | Thailand | $1.8 \times 10^6$ | IP | No | 71.3<sup>a</sup> | This study |
| | Formalin | | $1.8 \times 10^6$ | IP | No | 79.6<sup>a</sup> | |

601 IP, intraperitoneal injection; IM: intramuscular injection; a: Relative percentage survival (RPS); b: survival rate

### 4.2. Immunization with HKV or FKV activated both branches of the tilapia’s specific immune system

Upregulation in the expression of *IgM, IgD* and *IgT* and *CD4* (genes encoding proteins involved in humoral immunity) and *CD8* (cell-mediated immunity) following immunization with HKV and FKV suggests that the vaccines are able to activate both arms of the specific immune response in Nile tilapia. Protection from these vaccines is, therefore, likely to result from a synergistic effect of humoral (B cell) and cellular immune (T cell) responses. This is similar to the recent report by Zeng et al (2021a), showing that β-propiolactone-inactivated TiLV vaccines induced up-regulation of *MHC-I* and *MHC-II/CD4*, which belong to different arms of the immune system.

The increase in *CD4* transcripts at 14 and 21 dpv in fish vaccinated with HKV or FKV may reflect activated naïve CD4+ cells differentiating into helper T-cell subsets, Th1 and Th2. The Th1 cells produce cytokines that stimulate the expression of anti-viral and inflammatory genes, whereas cytokines secreted by Th2 cells stimulate the differentiation of B-cells into plasma cells to produce specific antibody (Secombes & Wang, 2012; Secombes & Belmonte; 2016; Smith et al., 2019). On the other hand, CD8 transcription was only seen to be significantly up-regulated in the spleen of the HKV group after booster vaccination, indicating that the HKV may stimulate CD8+ cell activation, which then differentiate into cytotoxic T-cells. These cells play a crucial role in cell-mediated immunity (Bo et al., 2012; Somamoto et al., 2002; Smith et al., 2019).

As well as assessing the expression of *IgM* transcripts, this study also examined the expression of two additional immunoglobulins IgD and IgT. Similar patterns of up-regulation were found in head kidney of fish after the primary immunization, suggesting that all three antibodies may be involved in the protective response elicited by the vaccines. Interestingly, significant increases in mRNA *IgT* levels were seen in the head kidney before booster vaccination and in the spleen after the booster vaccination for both the HKV and FKV groups, suggesting that IgT may be strongly associated with the protective response against TiLV. Unfortunately, the function of IgT in tilapia remains poorly understood. Functional localization studies in other fish species have shown that IgT plays an important role against infectious pathogens on mucosal surfaces, such as skin, gills and gut (Smith et al., 2019; Salinas et al., 2021; Zhang et al., 2011). Nevertheless, further studies are required to gain a better understanding on the role of IgT in tilapia’s defense system, especially in response to infection.

Although immune genes were significantly upregulated in the head kidney after primary immunization, this pattern of expression was not observed in the spleen. This suggest that the head kidney, apart from being a primary lymphoid organ, also act as an important secondary lymphoid organ where specific immune responses to the TiLV vaccine occurred. Studies in other fish have shown that the head kidney, containing blast cells, plasma cells and melano macrophages, is an important site for antigen presentation and antibody production (Kumar et al., 2016; Soulliere & Dixon, 2017). This might be similar in tilapia. However, it was unexpected to find no significant up-regulation of *IgM, IgT, IgD and CD4* in the head kidney at 7 days after the booster vaccination at 28 dpi. It is possible that the increase in gene expression occurred later than 7 days after the booster vaccination or in other secondary lymphoid organs (not assessed in this study). Therefore, future studies should investigate a longer time course for gene expression to better understand the dynamics of immune gene responses after booster vaccination.

### 4.3. HKV and FKV induce both systemic and mucosal IgM

In present study, HKV and FKV were shown to trigger both systemic and mucosal IgM responses, with similar patterns observed between the two vaccines. The increase in systemic and mucosal IgM in teleost is usually derived from the major lymphoid organs, such as head kidney and spleen (Zapata et al., 2006), but also from the mucosa-associated lymphoid organs located in the skin, gills, gut, or nasopharynx (not investigated in this study) (Smith et al., 2019; Salinas et al., 2021). In the present study, up-regulation of IgM expression occurred mainly in the head kidney, and to a less extent in the spleen, suggesting head kidney to be one of the main organs for IgM production in response to the TiLV vaccines. Although the pathway of IgM secretion in the mucosal compartment (mucus) is unclear, it is possible that mucosal antibodies are produced locally in the mucosa-associated lymphoid organs and/or by the systemic immune system (Esteban & Cerezuela, 2015; Koppang et al., 2015; Salinas et al., 2011; Salinas & Parra, 2015; Salinas et al., 2021). In other research using Asian seabass, monovalent and bivalent bacterial vaccines induced both systemic and mucosal IgM (Thu-Lan et al., 2021). Similar kinetics have been reported for IgM secretion in the serum of red hybrid tilapia, infected IP with TiLV (Tattiyapong et al., 2020). The levels of serum IgM increased significantly in Nile tilapia after immunization with β-propiolactone-inactivated virus (Zeng et al. 2021a) or with a recombinant vaccine based on segment 8 of TiLV (Zeng et al., 2021b). Mucosal IgM was not investigated in these studies, however. The presence of TiLV-specific IgM in the mucus of vaccinated fish suggests that these vaccines may be able to generate a primary immune response in multiple mucosal organs such as skin and gills, which are crucial sites to prevent the initial invasion of pathogenic agents (Esteban & Cerezuela, 2015; Koppang et al., 2015). The IgM levels produced by FKV was always slightly higher than HKV in both serum and mucus at all sampling points analyzed, indicating that FKV induces stronger systemic and mucosal IgM responses than HKV. This could be one of the factors explaining for slightly higher level of protection conferred by FKV.

In this study, increased levels of TiLV specific IgM after booster vaccination in both serum and mucus indicate successful induction of specific immune memory after first immunization. However, low levels of *IgM* mRNA detected at 28 dpv did not reflect the IgM levels measure by ELISA at this time point. It was likely that the earlier *IgM* transcripts had already degraded, while its translated products (antibody) remained. B-cells are the major component involved in humoral adaptive immunity. They are activated by specific antigen binding to the B-cell receptors on the cell, followed by presentation of processed antigens to naïve CD4-Tcells, which then differentiate into helper T-cells. With T cells’ help, B-cells differentiate into plasma cells and memory B-cells. Plasma cells are committed to antibody secretion, whereas memory B-cells are responsible for the long-lasting protection from subsequent exposure to the same pathogens (Secombes & Belmonte, 2016; Smith et al., 2019).

Although systemic and mucosal IgM levels were assessed in the study, we were unable to measure levels of other antibodies i.e. IgD and IgT by ELISA due to a lack of monoclonal antibodies for these immunoglobulin classes in tilapia. Further studies should investigate the cost of the vaccine for commercial production, the persistence of the immune response in vaccinated fish, duration of protection and efficacy testing these vaccines in a commercial setting.

In conclusion, this study reported on the efficacy of two simple TiLV inactivated vaccines without adjuvant (HKV and FKV) in preventing TiLV infection in Nile tilapia. The vaccines activated both branches of adaptive immunity, triggered expression of three immunoglobulin classes and elicited both systemic and mucosal IgM responses. Most importantly, these vaccines showed relatively high levels of protection against TiLV infection, and therefore seem very promising for the prevention of disease associated with TiLV.

## Acknowledgements

This study was financially supported by the GCRF Networks in Vaccines Research and Development, which was co-funded by the MRC and BBSRC and is supported by the International Veterinary Vaccinology Network. Thao Thu Mai acknowledges the PhD scholarship program of Chulalongkorn University for ASEAN and Non-ASEAN countries and the 90^th^ year anniversary of Chulalongkorn University Fund. The authors thank Ms. Putiya Sangpo and Mr. Dinh Hung Nguyen for their technical assistances.

## Data Availability Statement

The data that support the findings of this study are available on request.

## Author Contributions

Conceptualization, H.T.D., P.K., K.D.T.; investigation, T.T.M., S.T.; formal analysis, T.T.M., P.K., H.T.D.; methodology, S.S., S.T., J.Z.C, J.D.P, K.D.T, T.T.M., H.T.D; supervision, H.T.D., C.R.; writing - original draft, T.T.M, H.T.D., S.S.; review & editing, C.R., S.S., P.K., J.Z.C, J.D.P, K.D.T. All authors have read and agreed to the current version of the manuscript.

