## Supplementary material for "Efficacy of heat-killed and formalin-killed vaccines against *Tilapia tilapinevirus* in juvenile Nile tilapia (*Oreochromis niloticus*)": Table S1;Fig. S1

| Inactivated methods | Concentration (%) | Temperature (^o^C) | Time (hours) | CPE observation | | |
| --- | --- | --- | --- | --- | --- | --- |
|  |  |  |  | Rep.1 | Rep.2 | Rep.3 |
| Heat |  | 25 | 2 | + | + | + |
|  |  | 60 | 2 | - | - | - |
|  |  |  | 2.5 | - | - | - |
|  |  | 65 | 2 | - | - | - |
|  |  |  | 2.5 | - | - | - |
| Formaldehyde solution | 0 | 25 | 24 | + | + | + |
|  | 0.002 |  |  | - | - | - |
|  | 0.004 |  |  | - | - | - |
|  | 0.006 |  |  | - | - | - |
|  | 0.008 |  |  | - | - | - |
|  | 0.01 |  |  | - | - | - |

**Table S1.** Inactivation of TiLV using different methods

Rep, replicate; CPE, cytopathic effect; (+), have CPE; (-), no CPE.

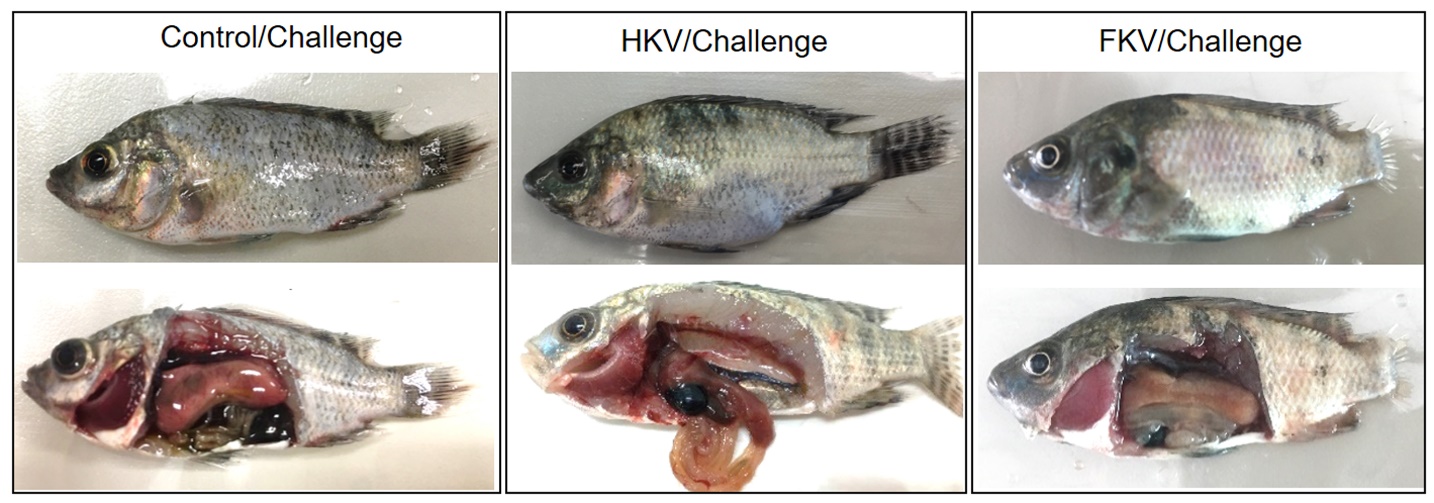

**Figure S1.** Gross signs of representative dead fish from control, HKV and FKH groups following challenge with TiLV. The fish showed common signs of TiLV infection, including abdominal distension, skin erosion, exophthalmos, fin rot, gill pallor and pale liver.
